# Transcriptional regulation of adipocyte lipolysis by IRF2BP2

**DOI:** 10.1101/2024.07.31.605689

**Authors:** Yang Chen, Lin Liu, Ryan Calhoun, Lan Cheng, David Merrick, David J. Steger, Patrick Seale

**Affiliations:** Institute for Diabetes, Obesity and Metabolism; Dept. of Cell and Developmental Biology, Perelman School of Medicine University of Pennsylvania

## Abstract

Adipocyte lipolysis controls systemic energy levels and metabolic homeostasis. Lipolysis is regulated by post-translational modifications of key lipolytic enzymes. However, less is known about the transcriptional mechanisms that regulate lipolysis. Here, we identify the transcriptional factor interferon regulatory factor-2 binding protein 2 (IRF2BP2) as a repressor of adipocyte lipolysis. Deletion of IRF2BP2 in primary human adipocytes increases lipolysis without affecting glucose uptake, whereas IRF2BP2 overexpression decreases lipolysis. RNA-seq and ChIP-seq analyses reveal that IRF2BP2 directly represses several lipolysis-related genes, including *LIPE* (*HSL*, hormone sensitive lipase), which encodes the rate-limiting enzyme in lipolysis. Adipocyte-selective deletion of *Irf2bp2* in mice increases *Lipe* expression and free fatty acid levels, resulting in elevated adipose tissue inflammation and glucose intolerance. Altogether, these findings demonstrate that IRF2BP2 restrains adipocyte lipolysis and opens new avenues to target lipolysis for the treatment of metabolic disease.

## Introduction

Adipocytes play a central role in regulating systemic energy levels and metabolic health. Adipocytes store energy as triglyceride in lipid droplets, and release energy in the form of free fatty acids (FFAs) through lipolysis ^1^. Adipocytes display remarkable plasticity and undergo dynamic changes in their metabolic program in response to many physiologic and pathologic stimuli.

Lipolysis is activated under conditions of energy demand, such as during fasting or exercise. Triglycerides are cleaved into diacylglycerol and fatty acids by the enzyme PNPLA2 (also called ATGL), followed by the hydrolysis of diacylglycerol into monoacylglycerol and fatty acids through the action of hormone-sensitive lipase E (LIPE, also known as HSL). Free fatty acids (FFA) are released into bloodstream to provide fuel for other organs. Excessive or dysregulated adipocyte lipolysis leads to ectopic fat deposition in liver, muscle, pancreas, and other organs, promoting insulin resistance and glucose intolerance ^2, 3, 4^. Aberrant lipolysis can also lead to hyperlipidemia and cardiometabolic abnormalities ^5, 6^. Mutations in key lipolysis genes (e.g. *PNPLA2*, *LIPE*) are associated with human metabolic and cardiac dysfunction, such as lipodystrophy, insulin resistance and cardiac myopathy ^7, 8^.

Adipocyte lipolysis is activated by the sympathetic nervous system and pro-lipolytic hormones such as cortisol, glucagon and natriuretic peptides ^4, 9, 10, 11^. These stimuli induce the phosphorylation and activation of lipolytic proteins PNPLA2 and LIPE ^12, 13, 14^. Conversely, insulin acts on adipocytes to restrain lipolysis ^11, 15, 16^. Adipocyte PNPLA2 and LIPE activities are also regulated via other various pathways e.g. cyclic adenosine monophosphate activators, FGF-1, serotonin, and inflammatory factors ^17, 18, 19, 20, 21, 22, 23^. However, there is limited research on the transcriptional regulation of lipolytic enzymes and lipolysis. Transcriptional mechanisms may be especially important for determining the rates of basal lipolysis. Peroxisome proliferator activated receptor gamma (PPARγ), liver X receptor alpha (LXRα), and steroidogenic factor-1 (SF1) have been shown to regulate *LIPE* expression ^24, 25, 26^. Identifying additional transcription factors and mechanisms that regulate lipolysis may provide new therapeutic avenues to ameliorate and/or prevent insulin resistance and related cardiometabolic disorders.

Interferon regulatory factor-2 binding protein-2 (IRF2BP2) is a transcriptional cofactor, governing diverse biological processes including macrophage inflammatory responses, lymphocyte differentiation, cardiomyocyte hypertrophy, and hepatic steatosis ^27, 28, 29, 30^. IRF2BP2 was first described as an IRF2-dependent transcriptional co-repressor but has also been reported to activate target genes in certain contexts ^31, 32^. Notably, *IRF2BP2* variants identified from genome wide association studies (GWAS) are associated with circulating lipid levels and coronary artery disease risk ^33^. However, the role of IRF2BP2 in regulating adipocyte function was unknown. In this study, we found that IRF2BP2 represses lipolysis in adipocytes, and this action is required to maintain systemic metabolic homeostasis. In human adipocytes, deletion of *IRF2BP2* elevates lipolysis, whereas overexpression of IRF2BP2 suppresses lipolysis. Integrated RNA-seq and ChIP-seq analysis demonstrate that IRF2BP2 binds and represses lipolysis-related genes, including the rate-limiting enzyme *LIPE*. Adipocyte-selective deletion of *Irf2bp2* in mice increases *Lipe* expression and leads to elevated levels of circulating FFA. Consequently, *Irf2bp2* deficient mice exhibit increased adipose tissue and systemic inflammation and glucose intolerance.

Altogether, our results demonstrate that adipocyte IRF2BP2 contributes importantly to whole body metabolic homeostasis through transcriptional repression of *LIPE* expression and lipolysis.

## Results

### Loss of *IRF2BP2* increases lipolysis in human adipocytes

We first evaluated the mRNA expression levels of *IRF2BP2* during human adipocyte differentiation. Primary human adipose tissue-derived precursor cells (hAPC) were induced to differentiate into lipid-droplet containing adipocytes that expressed high levels of adipocyte-selective genes (*ADIPOQ*, *FABP4*, *PPARG*, *LIPE*) by day 14 (**Extended Data Fig. 1a-c)**. *IRF2BP2* mRNA levels decreased by ∼50% at day 7 and remained lower in mature adipocytes compared to hAPC (**Fig. 1a**). Analysis of a single nucleus RNA-seq dataset from human adipose tissue also showed lower *IRF2BP2* mRNA levels in adipocytes relative to hAPC (**Extended Data Fig. 1d,e)** ^34^.

**Fig. 1:**
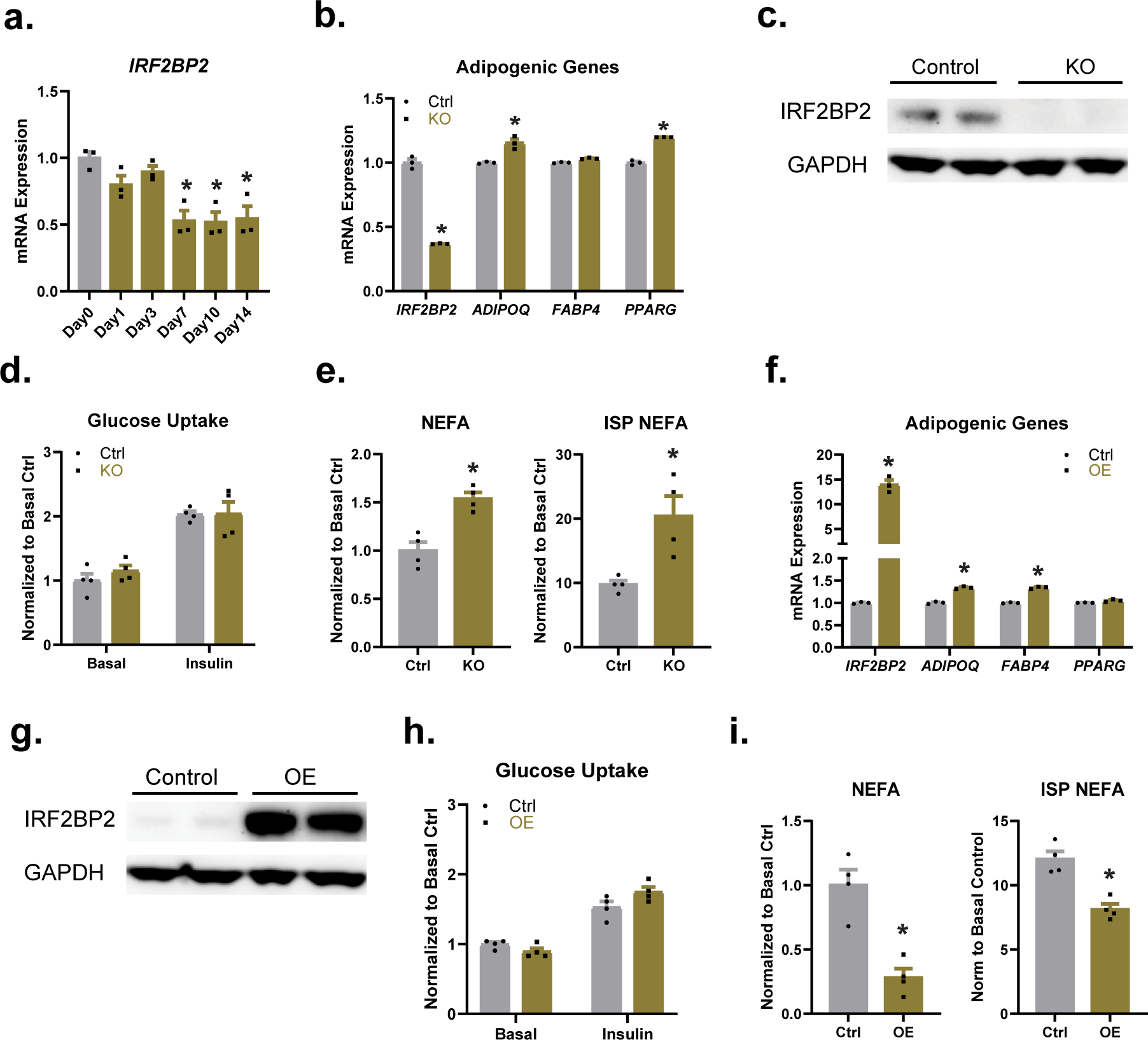
IRF2BP2 regulates adipocyte lipolysis. **a**, Relative IRF2BP2 mRNA levels during human adipocyte differentiation (n=3 in each time point). **b-e**, hAPC were transduced with IRF2BP2-targeting (knockout, KO) CRISPR lentivirus or non-targeting control lentivirus (Ctrl), and differentiated into adipocytes for 14 d. **b**, Relative mRNA levels of IRF2BP2 and adipocyte marker genes (*ADIPOQ*, *FABP4*, *PPARG*). **c**, Western blot analysis of IRF2BP2 and GAPDH (loading control) protein levels. **d**, Glucose uptake in Ctrl and KO adipocytes treated with either PBS or 10^-8^ M insulin. **e**, Non-esterified fatty acid (NEFA) levels in culture medium from Ctrl and KO adipocytes under basal conditions or following stimulation with 10^-6^ M isoproterenol (ISP). **f-i**, hAPC were transduced with IRF2BP2-expressing (overexpression, OE) or GFP-expressing lentivirus (Ctrl) and differentiated into adipocytes for 14 d. **f**, Relative mRNA levels of *IRF2BP2*, *ADIPOQ*, *FABP4*, and *PPARG*. **g**, Western blot analysis of IRF2BP2 and GAPDH (loading control) protein levels. **h**, Glucose uptake in Ctrl and OE adipocytes treated with either PBS or 10^-8^ M insulin. **i**, NEFA levels in culture medium from Ctrl and OE adipocytes under basal conditions or following stimulation with 10^-6^ M ISP. Unpaired two-tailed Student’s t tests were used in **b**, **e**, **f**, and **i**. One-way ANOVA followed by Dunnett multiple comparisons test was applied in **a**.

To evaluate the function of IRF2BP2 in human adipocytes, we deleted IRF2BP2 in hAPC using a lentiviral CRISPR system and induced adipocyte differentiation for 14 days. *IRF2BP2* mRNA levels were markedly reduced, and protein levels were near absent in *IRF2BP2*-knockout (KO) cells compared to control (transduced with non-targeting sgRNA) cells (**Fig. 1b,c**). Deletion of *IRF2BP2* had a minimal effect on the adipocyte differentiation process, with slight increases in the expression levels of adipocyte marker genes *ADIPOQ* and *PPARG* (**Fig. 1b**). *IRF2BP2* KO and control adipocytes (at day 14) also exhibited equivalent levels of basal and insulin-stimulated glucose uptake (**Fig. 1d)**. Interestingly, *IRF2BP2* KO adipocytes released significantly higher levels of non-esterified fatty acids (NEFA) into the culture medium under basal conditions, as compared to control cells (**Fig. 1e**). Isoproterenol treatment greatly increased NEFA release (lipolysis) in control and *IRF2BP2* KO adipocytes, with the KO adipocytes attaining higher levels (∼2-fold) of NEFA release compared to control cells.

### IRF2BP2 overexpression in human adipocytes decreases lipolysis

To determine if IRF2BP2 overexpression (OE) could reduce lipolysis, we transduced hAPC with control (GFP) or IRF2BP2-expressing lentivirus, and induced adipocyte differentiation for 14 days. IRF2BP2 OE cells expressed high levels of IRF2BP2 mRNA and protein and slightly increased the levels of adipocyte genes *ADIPOQ* and *FABP4* (**Fig. 1f,g**). Control and IRF2BP2-OE adipocytes displayed similar rates of glucose uptake under basal conditions and following insulin stimulation (**Fig. 1h**). Strikingly, IRF2BP2 expression decreased NEFA release by ∼70% under basal conditions, and significantly decreased NEFA levels following isoproterenol treatment (**Fig. 1i**). Taken together, these results demonstrate that IRF2BP2 represses NEFA release from human adipocytes without influencing the differentiation process or glucose uptake.

### IRF2BP2 represses lipolysis-related genes in adipocytes

To identify IRF2BP2-regulated genes and pathways in adipocytes, we performed bulk RNA-sequencing analysis of IRF2BP2-KO and IRF2BP2-OE adipocytes, along with their respective control adipocytes (**Fig. 2a,b**, **Extended Data Fig. 2a-f**). These studies identified high confidence sets of IRF2BP2-repressed genes (163 genes upregulated in KO and downregulated in OE cells) and IRF2BP2-activated genes (175 genes downregulated in KO and upregulated in OE cells). Pathway analysis of IRF2BP2-repressed genes identified “fatty acid oxidation” and “lipid catabolic process” as top-ranking terms (**Fig. 2c**, **Extended Data Fig. 2g**). Notably, the IRF2BP2-repressed gene set included several classical lipolysis-related genes e.g. *LIPE*, *HSD11B1*, *PNPLA2* (**Fig. 2e**). IRF2BP2-activated genes were enriched for pathways related to “cell migration”, “ossification”, and “extracellular matrix organization” (**Fig. 2d**, **Extended Data Fig. 2h**).

**Fig. 2:**
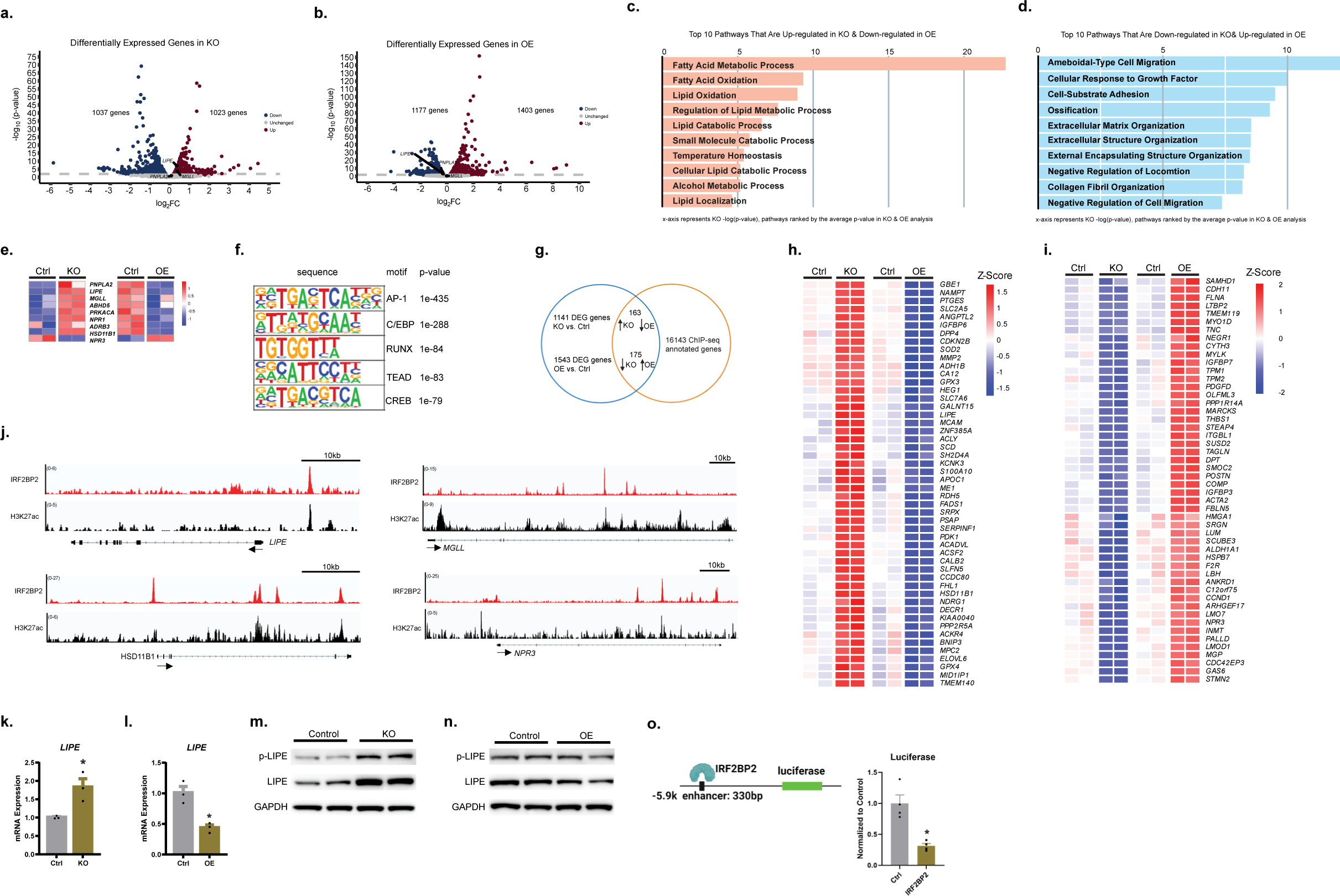
Identification of IRF2BP2 target genes including LIPE. **a-b**, Volcano plots showing results from RNA-seq analyses of control (Ctrl) vs. IRF2BP2-knockout (KO) (**a**) and Ctrl vs. IRF2BP2-overexpressing (OE) adipocytes (**b**). X axis: fold changes in mRNA levels (log2 transformed) over Ctrl group. Y axis: -log10 (p-value) for significance. **c-d**, Gene enrichment pathway analysis using Biological Processes (BP) database. **c**, The top 10 pathways that are: (1) upregulated in KO and (2) downregulated in OE adipocytes. **d**, The top 10 pathways that are: (1) downregulated in KO and (2) upregulated in OE adipocytes. **e**, Heatmap showing the expression profile of lipolytic genes in KO and OE adipocytes, relative to control cells. **f**, Motif analysis of IRF2BP2 binding regions in hAPC subjected to differentiation cocktail for 1 day. **g**, Venn diagram illustrates the overlap of genes identified from RNA-seq and ChIP-seq analyses. **h-i**, Expression heatmap of the top 50 IRF2BP2-repressed (**h**) and IRF2BP2-activated genes (**i**) that exhibit nearby IRF2BP2 binding peaks. **j**, ChIP-seq tracks for IRF2BP2 and H3K27Ac in *LIPE*, *MGLL*, *HSD11B1*, and *NPR3*. **k-l**, Relative LIPE mRNA levels in: (**k**) Ctrl and KO adipocytes and (**l**) Ctrl and OE adipocytes. **m-n**, Western blot analysis of LIPE, phospho-LIPE, and GAPDH (loading control) levels in: (**m**) Ctrl and KO adipocytes and (**n**) Ctrl and OE adipocytes. **o**, Transcription assay showing activity of -5.9 k region of *LIPE* in immortalized hAPC transfected with control vector (Ctrl) or IRF2BP2-expressing vector. For **k**,**l**,**o**, unpaired two-tailed Student’s t tests were used.

To identify genes that may be directly regulated by IRF2BP2, we performed chromatin immunoprecipitation sequencing (ChIP-seq) analysis for IRF2BP2 in hAPC following stimulation with adipogenesis cocktail. Motif analysis of IRF2BP2 binding regions identified a strong enrichment of motifs for several transcription factors, especially AP-1 and C/EBP (**Fig. 2f**). Integrated analysis of RNA-seq and ChIP-seq datasets was used to predict IRF2BP2 direct target genes by identifying genes that contain proximal binding sites and exhibit IRFBP2-regulated expression (repressed and activated, **Fig. 2g-i**). Several lipolysis-related genes (i.e. *LIPE*, *MGLL*, *HSD11B1* and *NPR3*) were identified as potential direct IRF2BP2-repressed genes (**Fig. 2h,j**). Many of these binding regions also displayed peaks of H3K27-acetylation, suggesting that they correspond to regulatory regions (**Fig. 2j**).

### IRF2BP2 directly represses *LIPE* transcription

We next focused on the IRF2BP2-regulation of *LIPE*, which encodes the rate-limiting enzyme in adipocyte lipolysis. qRT-PCR analysis showed that *LIPE* mRNA levels were increased (∼1.8 fold) by IRF2BP2 deletion and reduced (∼60%) by IRF2BP2 OE in human adipocytes (**Fig. 2k,l)**. LIPE protein levels were similarly regulated, with IRF2BP2-KO cells displaying increased LIPE protein levels and IRF2BP2-OE cells displaying lower LIPE levels compared to control cells (**Fig. 2m,n)**. Levels of phosphorylated (activated) LIPE followed a similar expression pattern (**Fig. 2m,n**). To determine if the IRF2BP2-binding region located 5.9 kb upstream of the *LIPE* transcriptional start site is regulated by IRF2BP2, we performed luciferase-based transcription assays in an immortalized hAPC cell line. IRF2BP2 expression decreased the transcriptional activity of this putative regulatory region by about 70%, relative to the vector control (**Fig. 2o**). These results suggest that IRF2BP2 represses *LIPE* transcription via binding to the -5.9 kb region.

### Adipocyte IRF2BP2 deficiency increases lipolysis and inflammation in mice

We next sought to determine the *in vivo* role of IRF2BP2 in adipocytes. To do this, we generated adipocyte-selective *Irf2bp2* knockout (AKO) mice by interbreeding *Irf2bp2^flox^* and *Adiponectin* (*Adipoq*)-*Cre* mice. *Irf2bp2* mRNA levels were significantly reduced in inguinal white adipose tissue (iWAT) and epididymal WAT (eWAT) of 12-week-old AKO compared to control mice (**Fig. 3a**). IRF2BP2 protein levels were also substantially reduced in WAT from AKO mice (**Fig. 3b**). Notably, *Irf2bp2* mRNA and protein levels were significantly higher in iWAT compared to eWAT (**Fig. 3a,b**). AKO and control mice had similar body weights at 12-weeks of age (**Fig. 3c**). iWAT weight was reduced by ∼25% in AKO mice compared to control mice, with no significant differences in either eWAT or brown adipose tissue (BAT) weights (**Fig. 3d,e**). H&E staining of iWAT sections showed that adipocyte size was reduced in iWAT of AKO compared to control mice, which likely accounts for the reduction in depot weight (**Fig. 3f,g)**. Circulating FFA was substantially elevated (∼75% higher) in AKO mice compared to control mice (**Fig. 3h)**. Consistent with the results in human adipocytes, *Lipe* expression was significantly elevated in iWAT (2-fold increased) and eWAT (1.5-fold increase) of AKO compared to control mice (**Fig. 3i)**. iWAT from AKO mice also expressed slightly higher levels of *Mgll* (**Fig. 3j)**. *Irf2bp2* deficiency did not affect the expression of *Adipoq*, or the lipogenic genes *Fasn*, *Scd*, and *Acly* (**Fig. 3k)**. Fasting for 16 hours reduced WAT weights and increased circulating NEFA levels to similar levels in control and AKO mice (**Extended data Fig. 3a,b**).

**Fig. 3:**
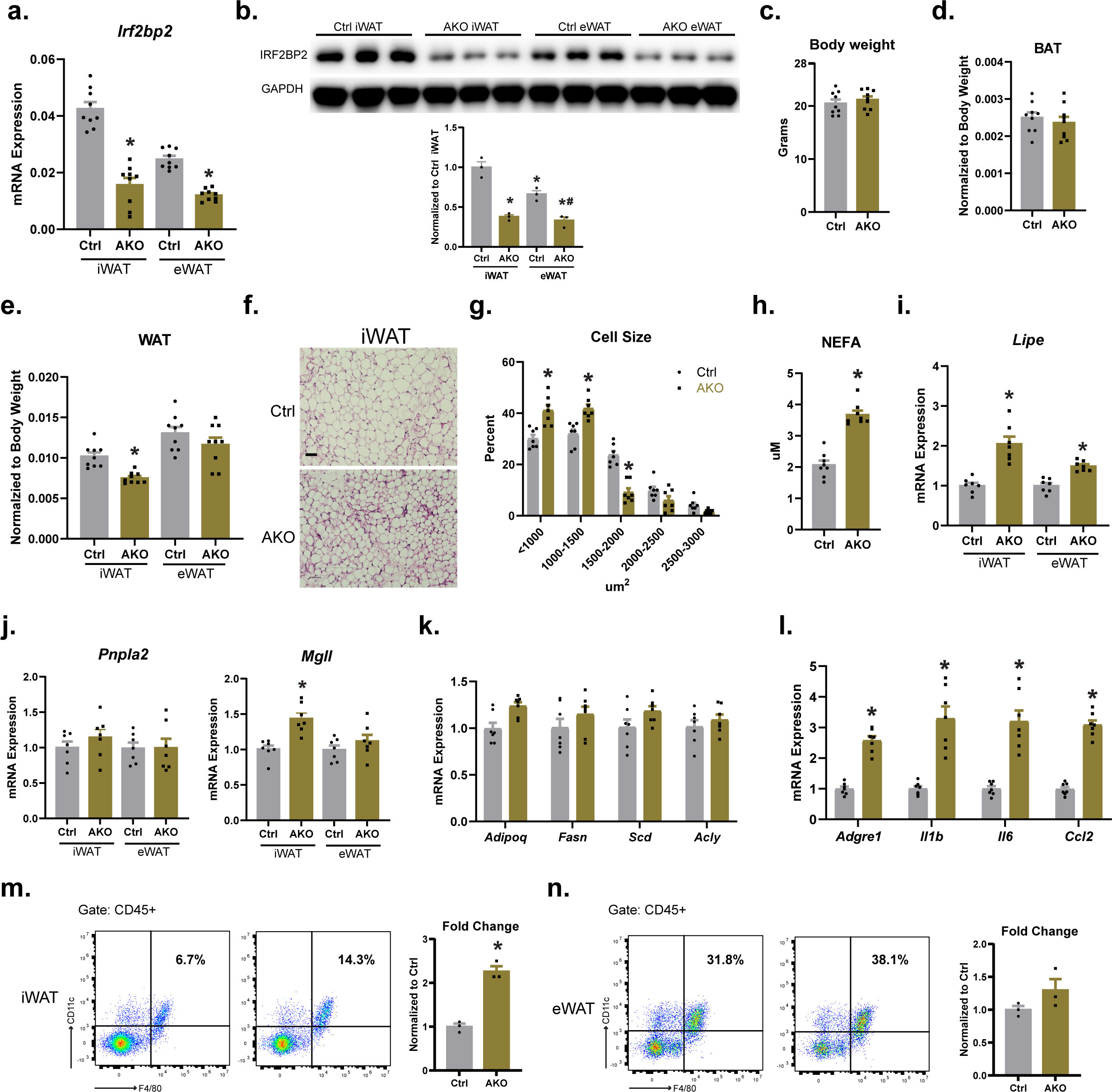
Adipocyte-specific deletion of Irf2bp2 increases LIPE expression, lipolysis, and inflammation in adipose tissue. Control and adipocyte-specific *Irf2bp2* knockout mice (AKO) were assessed as follows. **a**, Relative *Irf2bp2* mRNA levels in inguinal WAT (iWAT) and epididymal WAT (eWAT). **b**, Western blot analysis of IRF2BP2 and GAPDH (loading control) protein levels in iWAT and eWAT. Protein quantification was performed with imageJ. **c**, Body weights (n=9). **d-e**, Weights of: (**d**) brown adipose tissue (BAT), and (**e**) iWAT and eWAT (n=9). **f**, H&E staining of iWAT (scale bar, 50 um). **g**, Quantification of adipocyte cell size in iWAT (n=7). **h**, Circulating NEFA levels under ad libitum fed conditions (n=8). **i**, Relative Lipe mRNA levels in iWAT and eWAT (n=7). **j**, Relative mRNA levels of *Pnpla2* and *Mgll* in iWAT and eWAT (n=7). **k**, Relative mRNA levels of adipocyte genes *Adipoq*, *Fasn*, *Scd*, and *Acly* in iWAT (n=7). **l**, Relative mRNA levels of inflammatory genes *Adgre1* (F4/80), *Il1b*, *Il6*, and *Ccl2* (*Mcp1*) in iWAT (n=7). **m-n**, Flow cytometry analysis of F4/80+; CD11c+ (CD45+) cells in stromal vascular cells from iWAT (**m**) and eWAT (**n**) (n=3). In **a**, **b**, **e**, **h**, **i**, **j**, **l**, **m**, and **n**, unpaired two-tailed Student’s t tests were applied. Two-way ANOVA followed by Sidak’s test was applied for **g**.

Elevations in adipose tissue lipolysis and FFA are linked with macrophage recruitment, inflammation and metabolic dysfunction ^35, 36^. Consistent with this, we found that the expression of pro-inflammatory genes, including the macrophage marker *Adgre1* (i.e. *F4/80*), inflammatory cytokines *Il1b*, *Il6* and chemokine *Ccl2* were increased in iWAT of AKO versus control mice (**Fig. 3l**). Furthermore, flow cytometry analysis revealed a ∼2.5-fold increase in the proportion of pro-inflammatory F4/80^+^;CD11c^+^ cells in the stromal-vascular fraction of iWAT from AKO compared to control mice (**Fig. 3m**). The proportion of F4/80^+^/CD11c^+^ cells was also increased, but to a lesser extent, in the eWAT of AKO mice (**Fig. 3n**). Additionally, the circulating levels of inflammatory cytokines, IL6, IL1β, and MCP-1 trended to be higher in *Irf2bp2* AKO compared to control mice (**Extended Data Fig. 3c-e**). Hepatic triglyceride levels were increased in AKO mice, suggesting that adipose tissue-derived fatty acids accumulated in liver (**Extended Data Fig. 3f**). Together, these data indicate that IRF2BP2 represses adipocyte *Lipe* expression and suppresses lipolysis under basal conditions in mice.

### Adipocyte IRF2BP2 deficiency causes glucose intolerance in mice

We further tested if the action of IRF2BP2 in adipocytes regulates systemic glucose metabolism and insulin sensitivity. We found that fasting glucose levels were significantly elevated in AKO mice compared to control mice at 12 weeks of age (**Fig. 4a**). During a glucose tolerance test (GTT), *Irf2bp2* AKO mice exhibited higher blood glucose levels at each time point, with significant differences at 15 and 30 minutes (**Fig. 4b**). The area under the curve (AUC) for GTT was higher in *Irf2bp2* knockout mice (**Fig. 4c**). AKO mice also displayed reduced glucose-lowering effects of insulin during an insulin tolerance test (ITT), with elevated blood glucose levels at 0, 15, 60, 90, and 120 minutes (**Fig. 4d,e**). The data suggests that IRF2BP2-mediated suppression of basal lipolysis is required to maintain systemic glucose homeostasis.

**Fig. 4:**
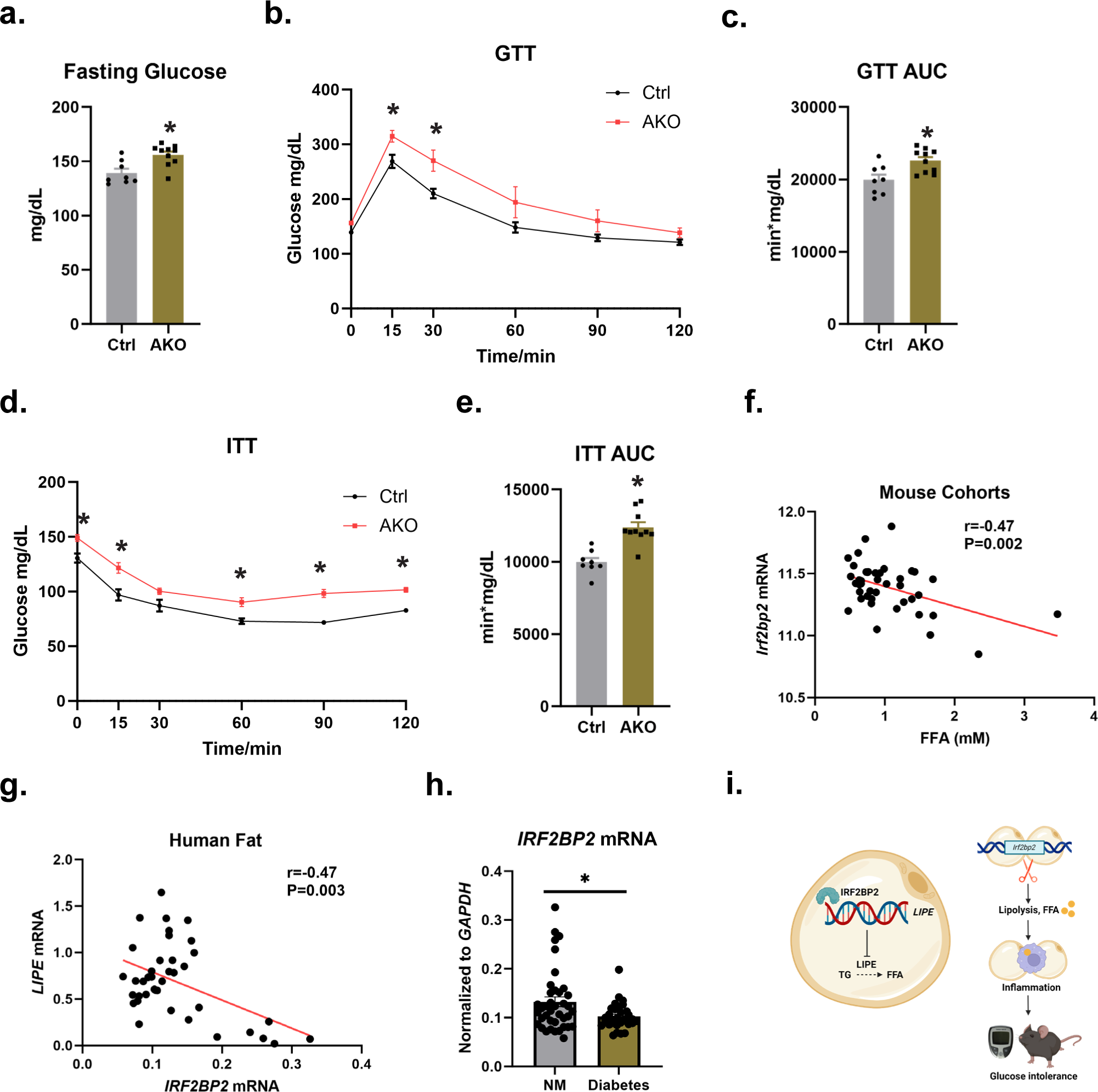
Adipocyte IRF2BP2 deficiency causes glucose intolerance. **a-d**, Analysis of metabolic parameters in 12-14 week-old control (Ctrl) and adipocyte-specific *Irf2bp2* knockout (AKO) mice (n=8-10). **a**, Fasting glucose levels. **b**, Intraperitoneal glucose tolerance test (GTT). **c**, GTT area under the curve (AUC) values from (**b**). **d**, Intraperitoneal insulin tolerance test (ITT). **e**, ITT AUC values from (**d**). f, Correlation between *Irf2bp2* mRNA levels in subcutaneous adipose tissue and serum free fatty acid (FFA) levels in 42 mouse strains (data extracted from GeneNetwork database, EPFL LISP3 Cohort) ^37^. **g**, Correlation between *IRF2BP2* and *LIPE* mRNA levels in subcutaneous adipose tissue from healthy, non-diabetic female subjects (n=39). **h**, *IRF2BP2* mRNA levels in subcutaneous adipose tissue from diabetic (n=31) and non-diabetic individuals (n=39). **i**, Working model: (left) IRF2BP2 transcriptionally represses *LIPE* expression in adipocytes to decrease the rate of lipolysis. (right) Loss of IRF2BP2 (or its downregulation in diabetes) increases lipolysis, leading to inflammation and impaired metabolic homeostasis. In **a**,**c**,**e**,**h**, unpaired two-sided t-tests were utilized. Two-way ANOVA followed by Sidak’s test was used in **b**,**d**. Person correlational analysis was performed in **f** and **g**.

### IRF2BP2 levels correlate with lipolysis markers in mice and humans

We next assessed if IRF2BP2 expression levels correlate with markers of lipolysis in mouse and human cohorts. In the GeneNetwork cohort consisting of 42 different normal mouse strains ^37^, *Irf2bp2* mRNA was significantly negatively correlated (r coefficient equals to -0.47) with circulating FFA levels (**Fig. 4f**). These data further support the role of IRF2BP2 as a negative regulator of adipocyte lipolysis in mice. To explore associations between IRF2BP2 and lipolysis in humans, we collected subcutaneous adipose tissue samples from non-diabetic (n=39) and diabetic subjects (n=31) for expression analysis by qRT-PCR. In the non-diabetic human cohort (n=39), *IRF2BP2* mRNA levels were negatively correlated (r=-0.47) with *LIPE* mRNA levels in subcutaneous adipose tissue (**Fig. 4g**). Interestingly, *IRF2BP2* mRNA levels were slightly (∼20%), but significantly decreased in adipose tissue from diabetic patients, compared to non-diabetic individuals (**Fig. 4h**). Taken together, these studies strongly support a conserved role of IRF2BP2 in restraining LIPE expression and lipolysis in mice and humans.

## Discussion

Dysregulated adipocyte lipolysis promotes the development of insulin resistance and cardiometabolic disease. In this study, we identified a novel role for the transcriptional cofactor IRF2BP2 as a repressor of adipocyte lipolysis. In human adipocytes, increased expression of IRF2BP2 decreased NEFA release, while loss of IRF2BP2 increased NEFA levels. In mice, deletion of IRF2BP2 in adipocytes elevated NEFA levels, and led to increased adipose tissue inflammation and systemic insulin resistance.

Our results suggest that IRF2BP2 restrains basal adipocyte lipolysis, but does not regulate the lipolytic response to catecholamines. In this regard, isoproterenol induced similar fold increases in lipolysis (i.e. NEFA levels) in IRF2BP2-expressing or IRF2BP2-deficient adipocytes relative to control adipocytes. Moreover, control and adipocyte-selective *Irf2bp2* knockout mice exhibited comparably high levels of circulating NEFA levels following a 16-hour fasting. Interestingly, the rate of basal (or resting) adipocyte lipolysis increases in obesity and correlates with cardiometabolic complications ^38, 39^. Additional studies indicate that lipolysis is increased in the early phase of impaired glucose tolerance in humans ^40, 41^. Mechanistically, fatty acids released by excessive lipolysis trigger pro-inflammatory cytokine production (i.e. IL-6, IL-1β, TNF-α) and provoke tissue inflammation ^42, 43^. Consistent with this, adipocyte-selective deletion of *Irf2bp2* led to adipose tissue inflammation and systemic insulin resistance in the absence of high fat diet or weight gain. These results suggest that chronic high levels of basal lipolysis are sufficient to trigger inflammation and insulin resistance.

Our results suggest that IRF2BP2 controls the rate of lipolysis via regulating the transcription of *LIPE*, which encodes the key lipolytic enzyme HSL. Previous studies revealed that upstream stimulatory factor (USF) and PPARγ bind to the *LIPE* promoter and modulate its transcription ^24, 25^. Our ChIP-seq analysis identified IRF2BP2 binds to a putative upstream enhancer region in the human *LIPE* gene. Previous studies suggest that IRF2BP2 functions as a transcriptional co-factor. Motif analysis of IRF2BP2 binding regions in adipocytes suggest that IRF2BP2 may interact with one or more member(s) of the AP-1 family of transcription factors e.g. ATF2, JUN FOS proteins to regulate *LIPE* enhancer activity. In this regard, IRF2BP2 was recently shown to interact and repress the function of the AP-1 heterodimer ATF7/JDP2 in leukemia cells ^44^. Studies employing JNK upstream kinase of JUN) and p38 (downstream effector of ATF2) inhibitors ^45, 46^ suggest that adipocyte lipolysis may be regulated by ATF2 and JUN, although the involved mechanisms remain unclear. Further studies are required to determine if IRF2BP2 interacts with AP-1 transcription factors to control LIPE expression and lipolysis in adipocytes. Moreover, our results show that IRF2BP2 regulates many other genes involved in lipolysis and adipocyte function, suggesting additional effects of IRF2BP2 beyond its regulation of LIPE levels.

Deletion of *Irf2bp2* had a more pronounced effect in the subcutaneous iWAT as compared to the visceral eWAT depot. It is well-established that iWAT and eWAT exhibit distinct responses to metabolic and inflammatory signals ^47^. eWAT undergoes higher rates of lipolysis ^47^ and harbors more pro-inflammatory macrophages ^48^. This may be related to the significantly higher expression levels of IRF2BP2 in iWAT. Thus, while IRF2BP2 disruption led to increased inflammation in both iWAT and eWAT, the effect was more consequential in iWAT due to its lower baseline levels. Future studies are warranted to explore the differential regulation and roles of IRF2BP2 in different fat depots.

Reducing fatty acid levels can ameliorate insulin action and improve cardiometabolic health, underscoring the therapeutic potential of targeting this pathway ^3, 5, 49, 50, 51^. For example, treatment of mice with the ATGL inhibitor, atglistatin, lowers circulating fatty acids, decreases adipose inflammation, reduces liver fat, and ameliorates glucose homeostasis ^50^. Similar findings are observed in *Mgll* mutant mice, *Lipe* haploinsufficient mice, and in mice treated with a LIPE inhibitor ^3, 51^. Increasing the function of IRF2BP2 in adipocytes may provide a selective strategy to blunt basal lipolysis and limit chronic inflammation for therapeutic benefit.

## Materials and Methods

### Mice

The mice used in our study were housed and cared for by the University of Pennsylvania University Laboratory Animal Resources (ULAR) in accordance with the guidelines set forth by the University of Pennsylvania Institutional Animal Care and Use Committee (IACUC). All animal procedures were performed following the guidance of the IACUC. Mice were maintained at room temperature on a normal chow diet and subjected to a 12-hour light/dark cycle unless specified otherwise. *Adipoq-Cre* mice were obtained from Jackson Laboratory (strain name: B6;FVB-Tg(Adipoq-cre)1dEvdr/J, RRID:IMSR_JAX:010803) ^52^. The *Irf2bp2* loxP/loxP strain was kindly provided by Dr. Alex Stewart ^27^.

Mice were euthanized by CO_2_ asphyxiation, followed by cervical dislocation. Blood (fed condition) was collected by cardiac puncture and subjected to centrifuge for plasma non-esterified fatty acids (NEFA) determination (HR series NEFA, WAKO). Fat tissues were immersed in 4% freshly prepared formaldehyde for histology analysis or immediately cryopreserved for protein and mRNA analysis.

### Mouse metabolic phenotyping

Glucose tolerance test (GTT) and insulin tolerance test (ITT) were conducted in male mice aged 12-14 weeks. For GTTs, a 6 h fasting period preceded the intraperitoneal (i.p.) administration of 1.5 g of glucose per kilogram, delivered as a 20% D-glucose solution dissolved in sterile water.

For ITTs, mice were fasted for 4 h, followed by a bolus i.p. injection of 0.65 U/kg insulin (Humulin, Novo Nordisk). Blood glucose levels were monitored from the tail-tip using an automatic glucometer (Contour Next, Bayer) at baseline and at 15, 30, 60, 90, and 120 minutes post-injection.

### Adipose tissue flow cytometry analysis

Stromal vascular cells were isolated from adipose tissue depots for analysis. Dissected and minced adipose tissue was washed in PBS followed by digestion in DMEM/F12 medium (Gibco Invitrogen) containing 2.4 U/ml of dispase, 1.5 U/mL of type I collagenase (Gibco Invitrogen), and 1% bovine serum albumin (BSA) at 37°C incubator with constant shaking for 30 minutes. The digestion was quenched with DMEM/F12 medium supplemented with 10% fetal bovine serum (FBS). The digested cell suspension was filtered through a 100 μm mesh filter (BD Biosciences) and centrifuged at 400 × g for 4 minutes at 4 °C. Red blood cells were lysed by incubating cell suspensions in a hemolysis buffer for 4 minutes. Cells were suspended in FBS-containing medium and further filtered using a 70 μm and a 40 μm mesh, followed by centrifugation (400 x g, 4 minutes). Cell pellets were re-suspended in FACS buffer (0.5% BSA, 2 mM EDTA in PBS) and incubated with fluorophore-conjugated antibodies for 45 minutes at 4°C in the dark. The following antibodies were used: CD45-APC/Cy7 (103116, BioLegend), F4/80-APC (123116, BioLegend), and CD11c-PE (12-0114-82, eBioscience). DAPI was used for viability staining. After three washes with FACS buffer, stained cells were suspended in 200 µL of FACS buffer for analysis. The CytoFlex Flow Cytometer (Beckman) was used to acquire 40,000 events for each sample. Unstained and single stain controls were used for setting compensation and gates. Debris, cell aggregation and dead cells (DAPI^+^) were excluded. Macrophages were identified as CD45^+^;F4/80^+^, and M1 macrophages were identified as CD45^+^;F4/80^+^;CD11c^+^. Data was analyzed using FlowJo v10 (Flowjo).

### Cytokine multiplex array assay

Mouse plasma samples were diluted 2-fold with PBS and subjected to cytokine multiplexing analysis using the Luminex 200 system (Luminex, Austin, TX, USA) by Eve Technologies Corp. We utilized the mouse Focused 10-Plex Discovery Assay (Millipore Sigma) according to the manufacturer’s protocol.

### Hepatic triglyceride measurement

Liver samples from male mice (age 12-14 weeks) weighing 50-100 mg were homogenized using a tissue lyser (25 Hz, 5 min) in a lysis buffer composed of 140 mM NaCl, 2.5% Triton-X, 0.2% sodium deoxycholate, and 50 mM Tris pH 7.4. Triglyceride levels were quantified employing the Stanbio Triglyceride LiquiColor kit. Following the manufacturer’s instructions, 2 µl of the sample or standard was added to 200 µl of the TG enzymatic mix, incubated for 10 minutes, and the reactions were subsequently assayed in a plate reader at 500 nm.

### Human adipose tissue-derived precursor cells (hAPC) and adipogenic differentiation

hAPC were obtained from subcutaneous adipose tissue samples collected through lipoaspiration. hAPC were differentiated into mature adipocytes using DMEM/F12 medium supplemented with 10% FBS and treated with a standard cocktail consisting of 2.5X10^-8^ M insulin, 10^-5^ M dexamethasone, 5X10^-4^ M IBMX, and 10^-6^ M rosiglitazone for 14 days. For ChIP and luciferase assays, hAPC were treated with the above differentiation cocktail without rosiglitazone.

### Lentivirus transduction

hAPC were seeded in 60 mm or 100 mm culture dishes and transduced with lentivirus using the lentivirus viraductin kit from Cell Biolabs. Lentivirus used in the experiments were produced by HEK239T cells (ATCC) in 150mm dishes. Lentivirus for *IRF2BP2* knockout was produced using lentiCRISPR v2 (Addgene plasmid # 52961) carrying guide RNAs (gRNAs) targeting *IRF2BP2*. The gRNAs were designed using CRISPOR ^53^ and IDT CRISPR web design tools and synthesized by IDT. IRF2BP2 OE lentivirus was produced using plenti backbone plasmid (Addgene plasmid # 17448) containing the full-length coding sequence cloned from hAPC. Plasmid sequences were confirmed by Sanger sequencing (Genewiz).

### qRT-PCR and RNA-seq analysis

Tissue RNA was harvested using Trizol (Invitrogen) according to the manufacturer’s protocol. Reverse transcription reactions were performed with High Capacity cDNA Reverse Transcription Kit (Thermo Fisher). Quantitative real time PCR (qRT-PCR) was performed using PowerUp SYBR Green Master Mix (Thermo Fisher Scientific) on a QuantStudio 5 instrument (Thermo Fisher Scientific). Primers used for qRT-PCR reactions were summarized in **Extended Table 1**. Relative gene expression levels were calculated by the ddCt method using the house-keeping gene *GADPH* (human) or *Gapdh* (mouse) as control.

For RNA-seq, extracted cellular total RNA samples were sequenced by NovaSeq 6000 PE150 (Novogene). The raw paired-end Fastq files were trimmed using the tool TrimGalore (version 0.6.5) with default parameters and paired option enabled. The trimmed files were then aligned to the GRCh38 (version 104) genome using kallisto (version 0.46.1) with the default parameters of the kallisto quant command for paired-end data ^54^. Kallisto output files were read into R (version 4.3.0) and differentially expressed genes were generated by EdgeR (version 4.0.2)^55^. Volcano plots were generated using the package ggplot2 (version 3.4.4) where genes with a p-value less than 0.05 were colored based on expression. The enrichment analysis was performed with gprofiler2 (version 0.2.2) using genes with a false discovery rate less than 0.1, positive or negative logFC cutoff, and being ordered by significance. A gSCS correction method was applied to the enriched terms, organism was set to hsapiens, ordered query to TRUE, a cutoff of 0.001 for significance, and using the "Biological Processes" as the source. The top enriched pathways were plotted into bar graphs using the ggplot2 package where the terms were ordered based on p-value for merged datasets. Heatmaps were generated using packages ComplexHeatmap (version 2.16.0) and pheatmap (version 1.0.12). Rows were scaled to their relative scores (Z-score) for both heatmaps. In **Extended Figure 2**, heatmaps rows were clustered in ComplexHeatmap by their Euclidean distances with a ward.D2 clustering method, and a k-means clustering of 4. The Venndiagram was generated using genes with a positive or negative foldchange and a false discovery rate cutoff of 0.1 in which genes were compared between the different datasets to evaluate overlapping differentially expressed genes across conditions for the RNA-seq.

### Lipolysis and glucose uptake in adipocytes

Fully differentiated adipocytes were serum starved overnight in DMEM/F12 medium. The cells were then treated with 2% BSA, serum-free medium containing either PBS (control) or 10^-6^ M isoproterenol for 4 h, and culture medium was collected for NEFA assessment (HR series NEFA, WAKO) following the manufacturer’s instructions. Briefly, 20 μl of the sample was added with 180 μl of solution A, followed by 60 ul of solution B. The absorbance was measured in a colorimetric plate reader at a wavelength of 550 nm. NEFA values were normalized to protein content, which was determined using Pierce BCA Protein assay. Glucose uptake was determined using a Promega Assay Kit (J1342). Briefly, overnight serum-starved adipocytes were washed with PBS, followed by treatment with 10^-8^ M insulin for 1 h in glucose-free medium. Cells were then subjected to 2-DG uptake, acid termination, neutralization, and luminescence measurement.

### Western blotting

Cells were lysed with RIPA lysis buffer (NaCl 150 mM, NP-40 1%, sodium deoxycholate 0.5%, SDS 0.1%, 50 mM Tris pH 7.4) containing protease inhibitors (Roche), phosphatase inhibitors (Thermo Fisher Scientific), and 100 uM PMSF (Sigma). Adipose tissue depots were homogenized in RIPA buffer using the Qiagen TissueLyser (28 hz, 4 min). Tissue homogenates were kept on ice for 30 min with intermittent vortex, followed by 2-3 rounds of centrifugation (15 min, 12000 g) to remove cell debris and lipids. Protein concentration was determined by BCA assay. 30 ug lysates were separated on precast 4-12% bis-tris NuPage gels (Thermo Fisher Scientific) and transferred to PVDF membranes. Proteins were detected using primary antibodies against IRF2BP2 (A303-190A, Bethyl Laboratories), phospho-LIPE (4126S, Cell Signaling, Danvers, MA), LIPE (4107S, Cell Signaling), and GAPDH (MA5-15738, Thermo Fisher Scientific). Horseradish-peroxidase-conjugated secondary antibodies i.e. anti-mouse (7076S, Cell Signaling) and anti-rabbit (7074S, Cell Signaling) were used, followed by SuperSignal West Pico PLUS ECL detection (Thermo Fisher Scientific). Blotting images were scanned using Amersham ImageQuant 800 (Cytiva). Bands were quantified using ImageJ.

### Dual-luciferase reporter assay

Luciferase reporter assay was performed as previously described ^56^. Immortalized hAPC were seeded in 48-well plates. Cells were treated with the differentiation cocktail of insulin, dexamethasone, and IBMX for 24 h, followed by transfection using lipofectamine 3000 method (Thermo Fisher Scientific). For 1 well, 0.75 ul lipo3000 and P3000 diluted with Opti-MEM medium (Thermo Fisher Scientific) were used. 200 ng of firefly luciferase enhancer reporter vector was co-transfected with 200 ng of IRF2BP2 or empty pcDNA3.1 vector, with 5 ng of PGK promoter-driven renilla luciferase as a normalization control for transfection efficiency. Cells were harvested 48 h post-transfection and lysates were subjected to luminescence measurement. Luciferase assay was performed using dual-luciferase reporter assay (E1910, Promega) according to the manufacturer’s instructions.

### ChIP and CUT & RUN sequencing

ChIP was performed as described previously ^57^. Cells were treated with the differentiation cocktail of insulin, dexamethasone, and IBMX for 24 h prior to chromatin collection. Sonicated input sample (10% of total chromatin) was saved and subjected to crosslink reversing and column purification. Sonicated chromatin samples were incubated overnight at 4°C with IRF2BP2 antibody in 1 ml ChIP buffer (50 mM HEPES pH 7.8, 140 mM NaCl, 1% Triton X-100, 0.1% Na-Deoxycholate and Complete protease inhibitor). IP was washed and then eluted by incubation at 65°C overnight in 50 μl ChIP elution buffer (50 mM Tris pH 7.5, 10 mM EDTA, 1% SDS) with 10 mM DTT. Beads were pelleted and the supernatant was transferred to a new tube. DNA was subsequently isolated by columns (NucleoSpin, Takara). H3K27ac CUT & RUN experiment was similarly performed in hAPC following manufacturer’s protocol (86652, Cell Signaling). DNA bound by H3K27Ac (Active Motif) or IgG (Millipore) was collected. ChIP and CUT& RUN libraries were prepared and sent for next-generation sequencing (Novogene). Sequencing data was analyzed as previously reported ^58^. Briefly, reads were aligned using Bowtie2 (version 2.5.0), followed by MACS2 (version 2.2.8) peak calling and Homer (version 4.11) annotation and motif enrichment.

### Histology

Tissues were fixed in 4% PFA overnight, washed in PBS, dehydrated in ethanol, paraffin-embedded and sectioned. Sections were stained with hematoxylin and eosin. Images were captured on a Keyence inverted microscope. Adipocyte size was quantified with ImageJ software.

### IRF2BP2 correlational studies in mice and humans

*Irf2bp2* mRNA transcript expression and free fatty acid (FFA) levels in 42 mouse strains were determined using data extracted from GeneNetwork database (i.e. EPFL LISP3 Cohort) ^37^. We assessed the mRNA levels of *IRF2BP2*, *LIPE*, and *GAPDH* (normalization control) in subcutaneous adipose tissue samples from non-diabetic (n=39) and diabetic (n=31) female subjects that underwent bariatric surgery with an approved IRB protocol at Hospital of the University of Pennsylvania.

### Statistical analysis

Data analysis was performed with GraphPad Prism 10 software. No power calculations were performed prior to initiation of the study. All individual data points were plotted to assay normality. Data normality was assessed with Shapiro-Wilk test. For normally distributed data, two-sided t tests were performed where comparisons between two groups were being assayed.

Nonparametric Mann-Whitney test was applied for data that were not normally distributed. Two-way and one-way ANOVA with pairwise comparisons were performed where comparisons between more than two groups were being assayed in cell or mice studies. Correlational analysis was performed using Graphpad.

## Acknowledgments

We are grateful for the *Irf2bp2* loxP/loxP mouse strain kindly provided by Dr. Alex Stewart from the University of Ottawa Heart Institute. We thank Dr. Jeff Ishibashi for his excellent technical support for CRISPR design.

## Author Contributions

L.L., R.C., L.C., and D.M. were involved in study design, conceived the idea, conducted the experiments, analyzed the data and drafted the paper; Y.C., D.S., and L.L. designed the study and performed the experiments; P.S. designed and supervised the study, obtained the funding and cowrote the paper.

