## supplemental figures/table for "Transcriptional regulation of adipocyte lipolysis by IRF2BP2"

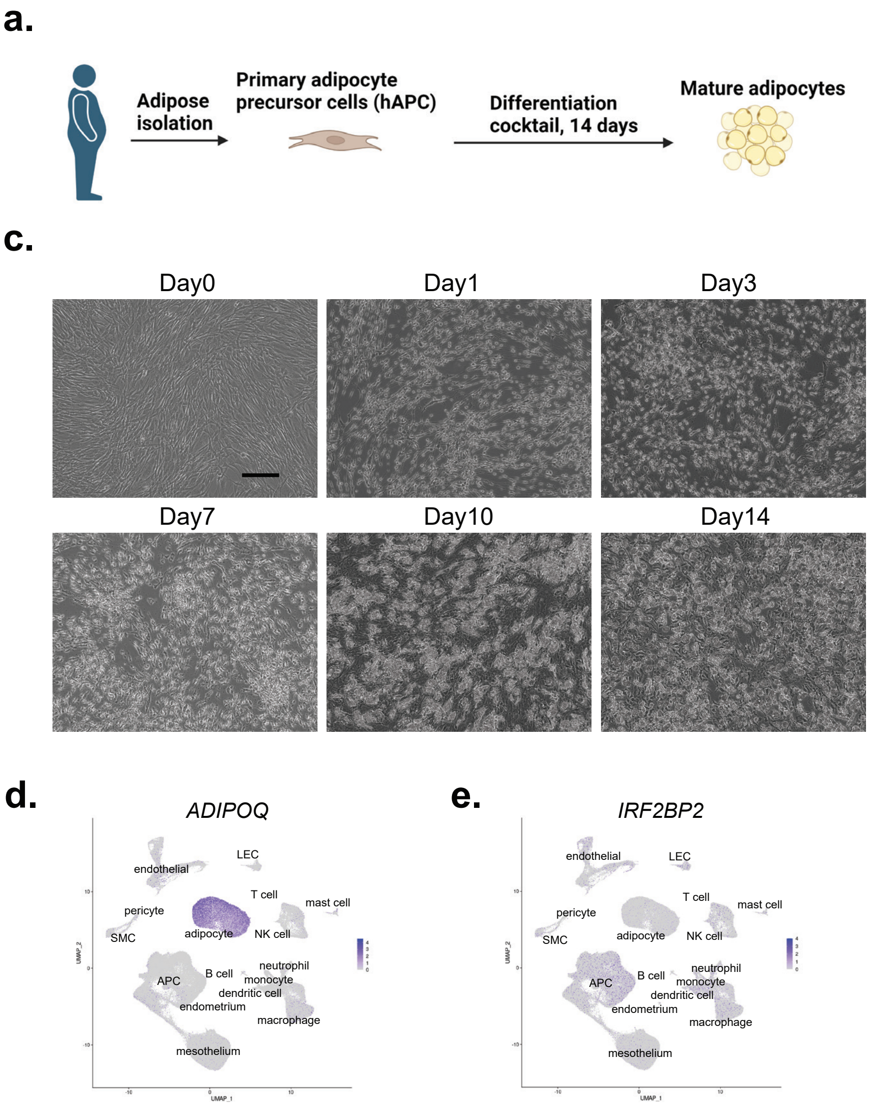


**Extended Data Fig. 1: *IRF2BP2* expression during human adipocyte differentiation**
**a**, Human adipocyte precursor cells (hAPC) isolated from subcutaneous adipose tissue were differentiated into mature adipocytes. n=3 per timepoint. **b**, Relative mRNA levels of adipocyte genes *ADIPOQ*, *FABP4*, *PPARG* and *LIPE* during differentiation. **c**, Phase contrast images of adipocyte cultures during differentiation (scale bar, 100 μm). **d-e**, UMAP of *ADIPOQ* and *IRF2BP2* expression in single nucleus RNA-seq dataset from human adipose tissue [34]. One-way ANOVA followed by Dunnett's test was used in **b**.

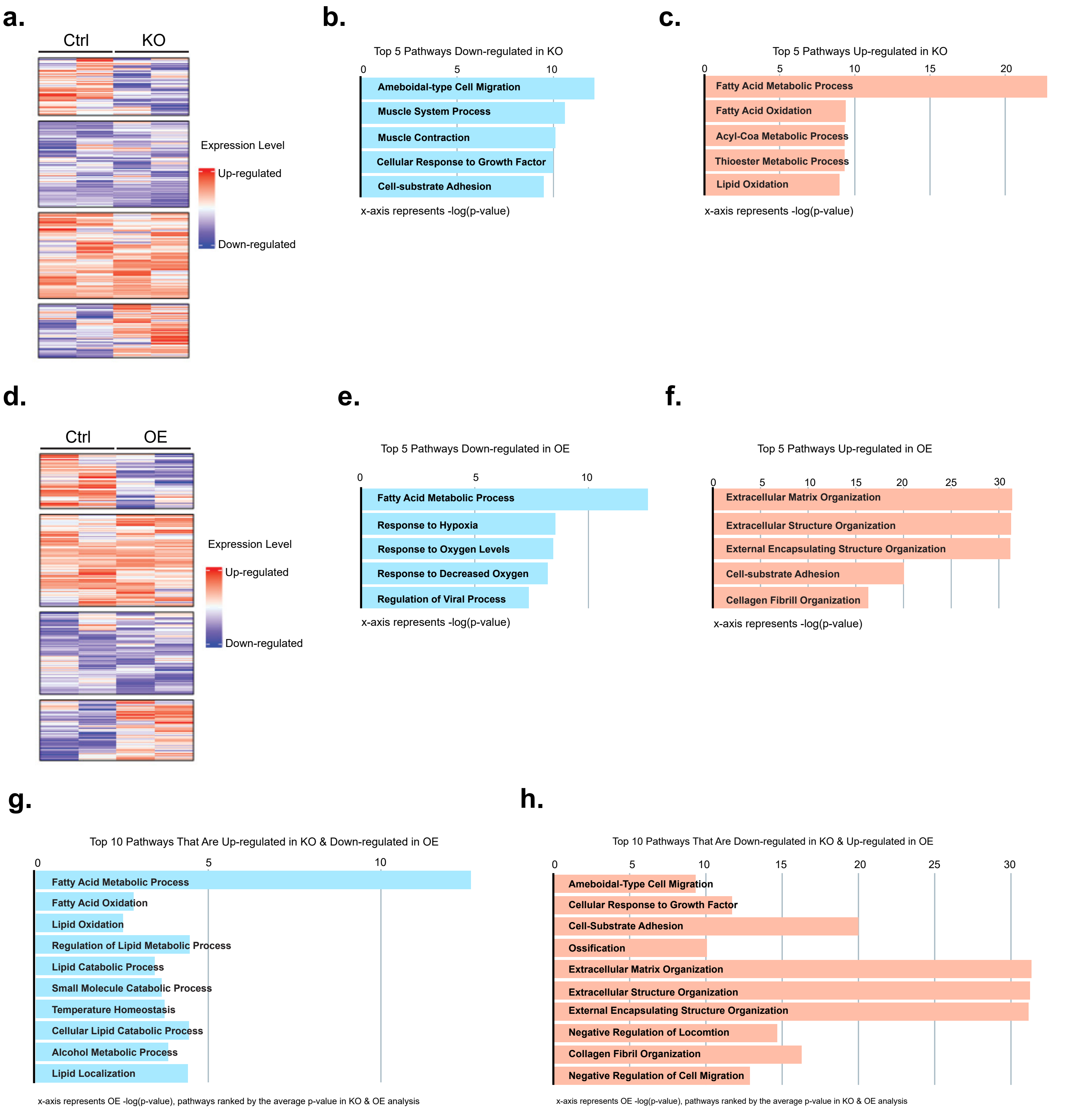


**Extended Data Fig. 2: Bulk RNA-seq heatmap and pathway analysis**
**a**, Expression heatmap from RNA-seq analysis of Ctrl and KO adipocytes. **b-c**, Biological process (BP) enrichment in Ctrl vs. KO cells: (**b**) downregulated pathways; and (**c**) upregulated pathways. **d**, Expression heatmap from RNA-seq analysis of Ctrl and OE adipocytes. **e-f**, BP enrichment in Ctrl vs. OE cells: (**e**) downregulated pathways; and (**f**) upregulated pathways. **g-h**, BP pathways that are upregulated in KO and downregulated OE, in Ctrl vs. OE study: (**g**) downregulated pathways; (**h**) upregulated pathways.

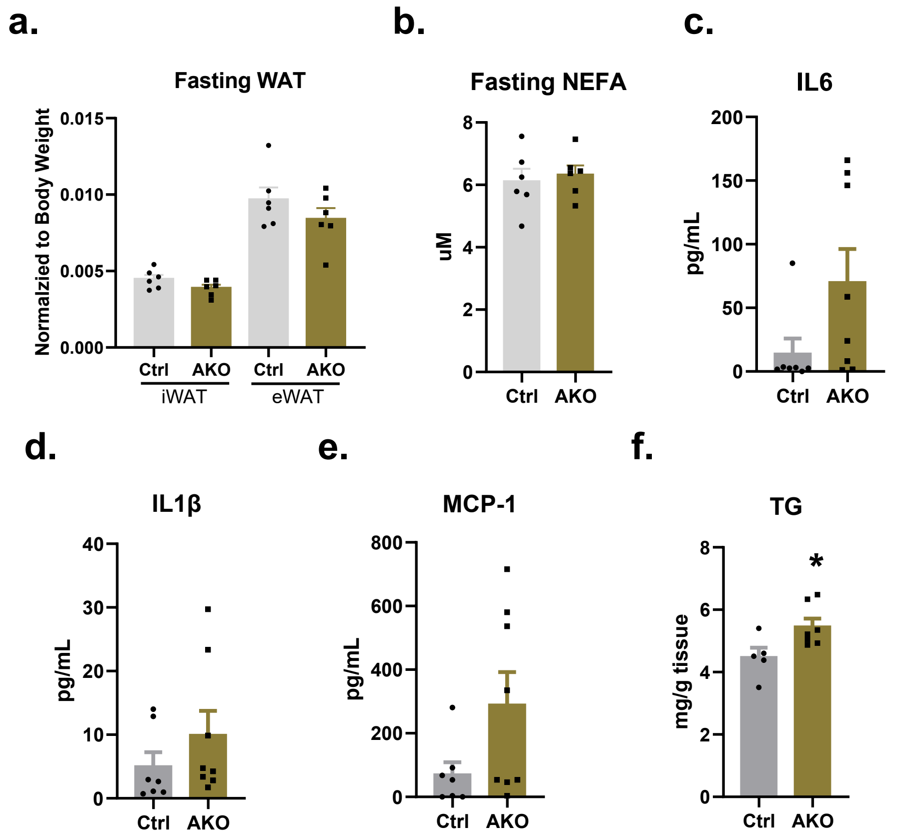


**Extended Data Fig. 3: Adipocyte IRF2BP2 regulates inflammation and metabolic parameters**
**a**, Adipose depot (iWAT, eWAT) weights in control and *Irf2bp2* AKO male mice (n=6, age 12-14 weeks) following a 16-hour fasting. **b**, Circulating NEFA levels in animals from (**a**). **c-e**, Circulating levels of: (**c**) IL6, (**d**) IL1β and (**e**) MCP-1 in 12-14 week old control and AKO male mice (n=7-8).

| Gene | Forward | Reverse |
|---|---|---|
| *GAPDH* | TCCAAAATCAAGTGGGGCGA | AAATGAGCCCCAGCCTTCTC |
| *IRF2BP2* | CATGATCTGGGACTTCACCG | TCTCGATGACGAACTCGACG |
| *PPARG* | AACAGATCCAGTGGTTGCAGA | CATGAGGGAGTTGGAAGGCT |
| *ADIPOQ* | AACATGCCCATTCGCTTTACC | TAGGCAAAGTAGTACAGCCCA |
| *FABP4* | CATGTGCAGAAATGGGATGG | AACTTCAGTCCAGGTCAACG |
| *LIPE* | AGTGCTTCTTCGCCTACTGC | GCAGATTCGTTCCCCTGTTG |
| *Gapdh* | GGCATTGTGGAAGGGCTCAT | AGATCCACGACGGACACATT |
| *Irf2bp2* | AGTTCTGTTTCCCTTGCTCC | TCTTCACTTTCACATCTCCGG |
| *Lipe* | GGGTGATGAAGGACTCACCG | GATGGCAGGTGTGAACTGGA |
| *Adgre1* | CTCAGTCTGCACCAATATCCTG | CCACAGAGTTAGAGCAGTTGGAA |
| *Il1b* | GTGTCTTTCCCGTGGACCTT | AATGGGAACGTCACACACCA |
| *Il6* | AGAGACTTCCATCCAGTTGCC | CCGGACTTGTGAAGTAGGGAA |
| *Ccl2* | AGGTGTCCCAAAGAAGCTGT | AAGACCTTAGGGCAGATGCAG |
| *Plnla2* | TTCGCAATCTCTACCGCCTC | AGCAAAGGGTTGGGTTGGTT |
| *Mgll* | CGCGCAGTAGTCTGGCTCTA | ATTCTGTGGAGTTCGCCTGG |
| *Adipoq* | ATCTGGAGGTGGGAGACCAA | GGGCTATGGGTAGTTGCAGT |
| *Fasn* | GCTGCGGAAACTTCAGGAAA | GAGTTGAGCTGGGTTAGGGT |
| *Scd* | CGAGGGCTTCCACAACTACC | AACTCAGAAGCCCAAAGCTCA |
| *Acly* | ACCATCATTGGGCCAGCTAC | GACATGCCTCCTGAACGTGA |

**Extended Table 1: qRT-PCR primer sequences**
